# The Greenland shark genome: insights into deep-sea ecology and lifespan extremes

**DOI:** 10.1101/2025.02.19.638963

**Authors:** Kaiqiao Yang, Kazuya Nishiwaki, Hideaki Mizobata, Shuichi Asakawa, Kazutoshi Yoshitake, Yuuki Y. Watanabe, Nigel E. Hussey, Kit M. Kovacs, Christian Lydersen, Mitsutaka Kadota, Shigehiro Kuraku, Shigeharu Kinoshita

## Abstract

The Greenland shark (*Somniosus microcephalus*) is known for its slow metabolism and deep-sea habitat. It is considered the longest-lived vertebrate on Earth, with an estimated lifespan of 392±120 years. Despite its remarkable longevity and lifestyle, there have been no genomic studies on this species. Here, we report the first, chromosome-level assembly of the Greenland shark genome, which is 5.9 Gb in size with an N50 length of 233 Mb, and contains 37,125 predicted genes with a completeness score of 86.5%. We found that the copy numbers of three gene families (*TNF*, *TLR*, *LRRFIP*), which are involved in activating the NF-κB signaling pathway, are significantly increased in the Greenland shark compared to short-lived shark species. In the rhodopsin of this deep-sea dweller, we detected amino acid substitutions that result in spectral tuning for the so-called ‘blue shift’, suggesting adaptive evolution to dim-light vision. We also elucidate the dynamics of the effective population size (*N_e_*) of the Greenland shark and its close relative, the Pacific sleeper shark (*Somniosus pacificus*). These genomic analyses offer new insights into the molecular basis of the exceptional longevity of the Greenland shark and highlight potential genetic mechanisms that could inform future research into longevity.

## Introduction

Longevity has been the subject of vast amounts of research effort. Studies on classical aging model organisms, such as mice, fruit flies, and zebrafish^1^, have uncovered key hallmarks of aging, including genomic instability, telomere attrition, epigenetic alterations, and loss of proteostasis^2^. Although aging appears to be inevitable, some animals have achieved exceptional longevity. Given the recent decline in sequencing costs and advances in bioinformatics, comparative genomic studies of long-lived species have become increasingly available, revealing unique alterations in various aging-related genes. Examples include the increased copy number of the *TP53* gene in the elephant genome^3^, the expansion of the immune modulatory butyrophilin gene family in the rockfish genome^4^, and specific sequence variations in the growth hormone receptor and insulin-like growth factor 1 receptor in Brandt’s bat genome^5^.

The Greenland shark (*Somniosus microcephalus*) is a large Squaliforme, which inhabits deep waters in the North Atlantic and Arctic Oceans. It is one of the largest extant shark species, reaching lengths >6 meters and weighing up to 1,400 kilograms, with a very slow annual growth rate (1 cm/yr)^6,7^. This species is known to undertake complex long-distance movements, with biotelemetry studies indicating seasonal inshore-offshore shifts^8^, multiyear returns to inshore sites^9^, and large-scale migrations that exceed 1000 km^10–12^, despite being one of the slowest swimming fish in the ocean^13^. Stomach content analyses show they prey on a variety of marine animals, including seals^11^, and they are considered a top predator as well as a scavenger in the Arctic Ocean^14^, although their corneas are frequently parasitized by *Ommatokoita elongata*, which was believed to impair their vision^7^. In 2016, age estimation of *S. microcephalus* using radiocarbon dating of the eye lens revealed that one individual was 392±120 years old^15^. This suggests that *S. microcephalus* is the longest-lived vertebrate, with a lifespan that surpasses even that of giant tortoises^16^ and the bowhead whale^17^.

Due to the technical challenges posed by large genome sizes and the abundance of repetitive sequences, it has proven to be challenging to unravel the diversity in morphology, reproductive mode, and lifespan of elasmobranchs. Since the first reported genome assembly for a chondrichthyan species, the elephant shark (*Callorhinchus milii*) in 2014^18^, genome assemblies were created for only the whale shark (*Rhincodon typus*)^19^ and the white shark (*Carcharodon carcharias*)^20^ in the following years. In 2020, the Squalomix consortium was established to provide genomic data, including genome sequences, transcriptomes, and epigenomes, in the hope of improving the global infrastructure for chondrichthyan genomic research^21^. Such work is important for conservation and international shark management, it is known that extant chondrichthyan species have been heavily impacted during the Anthropocene^22^ with global declines in populations reported^23^, but data limitations have inhibited appropriate fisheries and conservation management^24^.

*S. microcephalus*, which is often caught as bycatch in commercial fisheries, is estimated to take 150 years to reach maturity^15^. Such highly k-selected life history traits would suggest this species cannot withstand high levels of exploitation over long periods^25^. As a result, it was recently listed as a vulnerable species on the IUCN Red List in 2019 under criteria A2bd^26^. Consequently, obtaining genomic information for such species is increasingly seen as an important tool for understanding evolutionary processes but also to assess population dynamics to mitigate biodiversity loss. However, the genomic resources for the *S. microcephalus* are currently limited to the mitochondrial genome^27^, and the genetic basis for the species’ longevity remains unclear.

To alleviate some existing data gaps, we sequenced the whole genome of *S. microcephalus* and report herein the first chromosome-level genome for the species. Through population genetics analyses of *S. microcephalus*, and comparative analyses with other chondrichthyan genomes, we reveal genetic traits associated with cancer, immune response function, genome stability and cardiac function, deep-sea adaptations, and population size dynamics.

## Results

### High completeness genome assembly of *S. microcephalus*

Our high-fidelity long-read genome sequencing yielded approximately 34.5x coverage of the whole genome. By scaffolding the contigs we obtained 42 chromosome-scale sequences (Fig. 1B). The final genome assembly consisted of 5.9 Gb, with an N50 length of 233 Mb. The QV and k-mer completeness scores were 61.0 and 92.8, respectively (Table 1). Compared to the results of other large genome assemblies, such as the TrioCanu human (NA12878) assembly tested in the Merqury study with a QV score of 31^28^, our results appear to be very accurate. Comparisons with the genome assemblies of three other shark species with relatively large genome sizes reveal a markedly longer N50 length, as well as modest quantities of undetermined regions and non-chromosomal sequences (Fig. 1D) in *S. microcephalus*. High coverage of the protein-coding landscape estimated by single-copy ortholog retrieval (96.7%) also shows the high continuity and completeness of our assembly, despite its large size (Fig. 1D).

**Figure 1.**
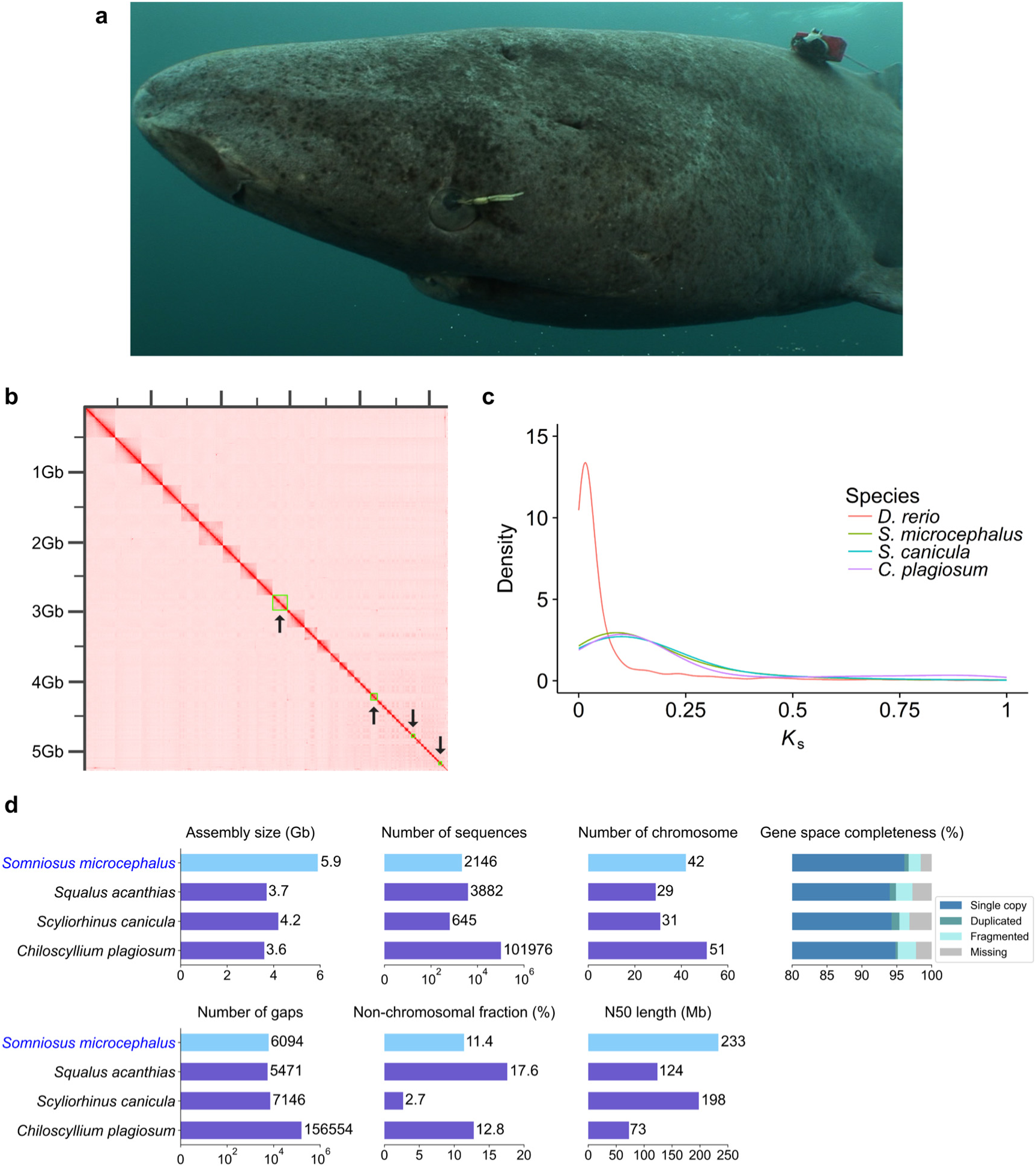
Assembling the *S. microcephalus* genome. (A) *S. microcephalus* (B) Hi-C heatmap of chromatin interaction counts, with boxes and arrows indicating pseudochromosomes 10, 20, 30, and 40. (C) Paralogs *K*_s_ distribution plots for three chondrichthyan species and *D. rerio*. (D) Comparison of genome assembly statistics between *S. microcephalus*, *S. acanthias*, *S. canicula*, and *C. plagiosum*.

**Table 1.** Statistics of *S. microcephalus* genome assembly.

| Assembly feature | Statistics |
| --- | --- |
| Assembled genome size | 5.9 Gb |
| Contig N50 | 1.64 Mb |
| Number of contigs | 9,670 |
| Scaffold N50 | 233 Mb |
| Number of scaffolds | 2,136 |
| Longest scaffold | 431 Mb |
| GC content of the genome | 47.77% |
| Completeness score | 96.7% |
| QV scores | 61.0 |
| k-mer completeness | 92.8% |
| Coverage | 99.9% |
| Mean depth | 34.5 |

Sequence determination of the *Cytb* region confirmed that the individual in this study possesses the H18 haplotype of *S. microcephalus* (EF090946.1)^29^, with 100% identity in a 703 bp-long stretch. Sequence determination of the *RAG1* and *ITS2* regions also confirmed that this individual possessed the *S. microcephalus*-specific sequences, with matches of 659/659 bases for *RAG1* (MF555591.1) and 1085/1086 bases for *ITS2* (MF555593.1)^30^, with no signs of hybridization.

In total, we annotated 4.97 Gb of repetitive sequences, representing 83.1% of the whole genome. Among these, retroelements, DNA transposons, and tandem repeats accounted for 46.39%, 3.88%, and 12.9% of the whole genome, respectively (Table S1). We obtained 37,125 protein-coding genes with a completeness score of 86.5% (Table S2). Searching the predicted gene models against publicly available databases including eggNOG, InterPro, UniProt, NR, GO, KEGG, etc., we functionally annotated 32,132 protein-coding genes, representing 86.5% of all protein-coding genes (Table S3). We also predicted genomic regions for 13 non-coding RNA types, including 15,563 transfer RNAs (tRNAs), 906 ribosomal RNAs (rRNAs), 241 microRNAs (miRNAs), and 1,221 small noncoding RNAs (snRNAs) in the genome assembly (Table S4). Among the rRNA, 816 ribozymes are located within 42 pseudochromosomes, comprising 90% of the total, highlighting the high quality and completeness of the genome assembly.

### Unique gene repertoires of *S. microcephalus* implicated in its exceptional lifestyle

To explore the evolutionary history of *S. microcephalus* and the potential biological role of expanded and contracted gene families that might facilitate longevity, we used the *S. microcephalus* genes and those of 12 chondrichthyan species (*Amblyraja radiata, Leucoraja erinacea, Pristis pectinata, Mobula hypostoma, Hypanus sabinus, Squalus acanthias, Hemiscyllium ocellatum, Chiloscyllium plagiosum, Rhincodon typus, Stegostoma tigrinum, Scyliorhinus canicula, Callorhinchus milii*) as well as an outgroup (*Danio rerio*) for orthology analyses (Table S5). OrthoFinder^31^ assigned 556,369 genes (95.8% of the total) to 25,463 orthogroups. Among these, 9,666 orthogroups contained all species, with 371 consisting entirely of single-copy genes. Additionally, 454 gene families were unique to *S. microcephalus* (Supplementary File 1); GO and KEGG enrichment results are shown in Figures S2 and S3. Based on the 371 single-copy genes, our reconstructed phylogenetic tree achieved full bootstrap support. Molecular dating using two fossil records suggests that *S. microcephalus* and *S. acanthias* diverged approximately 148.5 million years ago (Fig. 2A).

**Figure 2.**
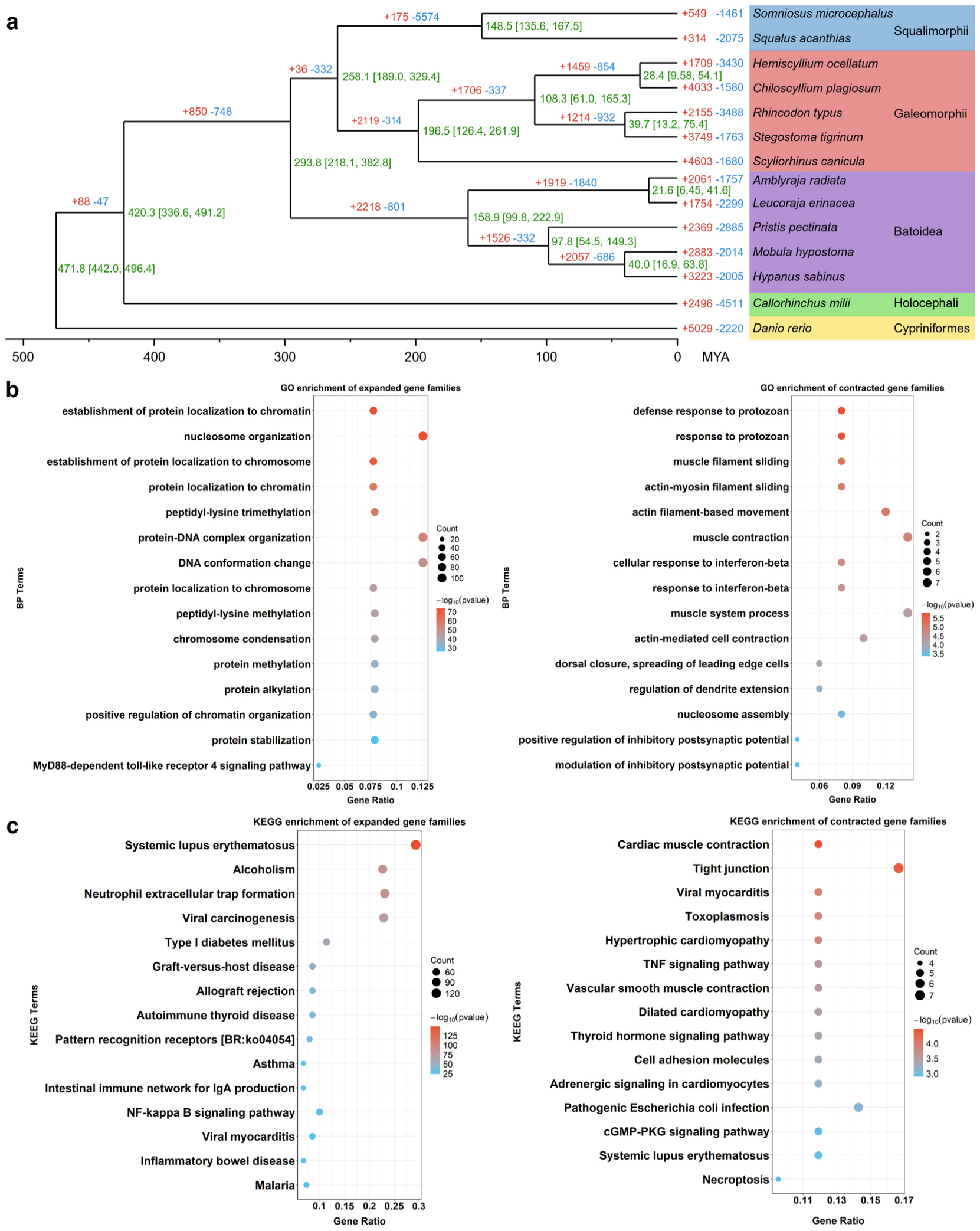
Analysis of gene family expansion and contraction. (A) Phylogenetic tree of 14 fish species. Green numbers on nodes represent estimated divergence times, with the 95% highest posterior density (HPD) in brackets. Red and blue numbers represent expanded and contracted gene families in each species and its common ancestor. (B) Top fifteen GO biological process terms of significantly expanded and contracted gene families. (C) Top fifteen KEGG enrichment terms of significantly expanded and contracted gene families.

To determine whether whole-genome duplication (WGD) events occurred, we calculated the *K*_s_ distributions of paralogous genes for each of *S. microcephalus* and three other species. The result indicates that *S. microcephalus* shares a *K*_s_ peak with two other chondrichthyan species at a larger x-axis value, while *D. rerio* exhibits a *K*_s_ peak at a smaller x-axis value specific to its lineage (Fig. 1C).

We identified 549 expanded and 1,461 contracted gene families in *S. microcephalus* (Fig. 2A). Using a p-value threshold of 0.05, 66 gene families were significantly expanded and 56 were significantly contracted (Supplementary File 2). The top 20 terms for GO and KEGG enrichment of significantly expanded and contracted gene families are shown in Fig. 2B and 2C, with the complete results in Supplementary File 3. GO enrichment analysis of expanded gene families identified numerous biological process terms related to chromosomal stability and DNA repair, such as positive regulation of chromatin organization, chromosome condensation, DNA conformation change, and protein-DNA complex organization. Additionally, terms associated with cell growth and division, such as positive regulation of cell growth and female meiosis chromosome segregation, were identified. GO enrichment analysis of contracted gene families revealed terms related to muscle function, including muscle filament sliding, actin-myosin filament sliding, and muscle contraction. KEGG enrichment analysis identified numerous biofunctional pathways, including those involved in autoimmune diseases (e.g., systemic lupus erythematosus, type I diabetes mellitus, autoimmune thyroid disease, rheumatoid arthritis), infectious diseases (e.g., viral carcinogenesis, viral myocarditis, malaria, *Staphylococcus aureus* infection, toxoplasmosis, pathogenic *Escherichia coli* infection), and cardiovascular-related pathways (e.g., vascular smooth muscle contraction, dilated cardiomyopathy, adrenergic signaling).

We found a significant expansion of genes associated with the NF-κB signaling pathway in *S. microcephalus* compared to other shark species (Fig. 3A). We examined the gene copy numbers for the *TNF* (tumor Necrosis Factor), *LRRFIP* (leucine-Rich Repeat Flightless-Interacting Protein), and *TLR* (toll-Like Receptors) gene families in five shark species and confirmed the significant expansion of all three gene families in *S. microcephalus* (Fig. 3B). The *TNF* family, comprising *TNFα* and *TNFβ*, the *LRRFIP* gene family, including *LRRFIP1* and *LRRFIP2*, and the *TLR* gene family have 14, 12, and 31 copies, respectively, in *S. microcephalus* (Fig. 3C). Additionally, the *CGNL1* gene, which is associated with cell junctions, has expanded to 14 copies (Supplementary File 2).

**Figure 3.**
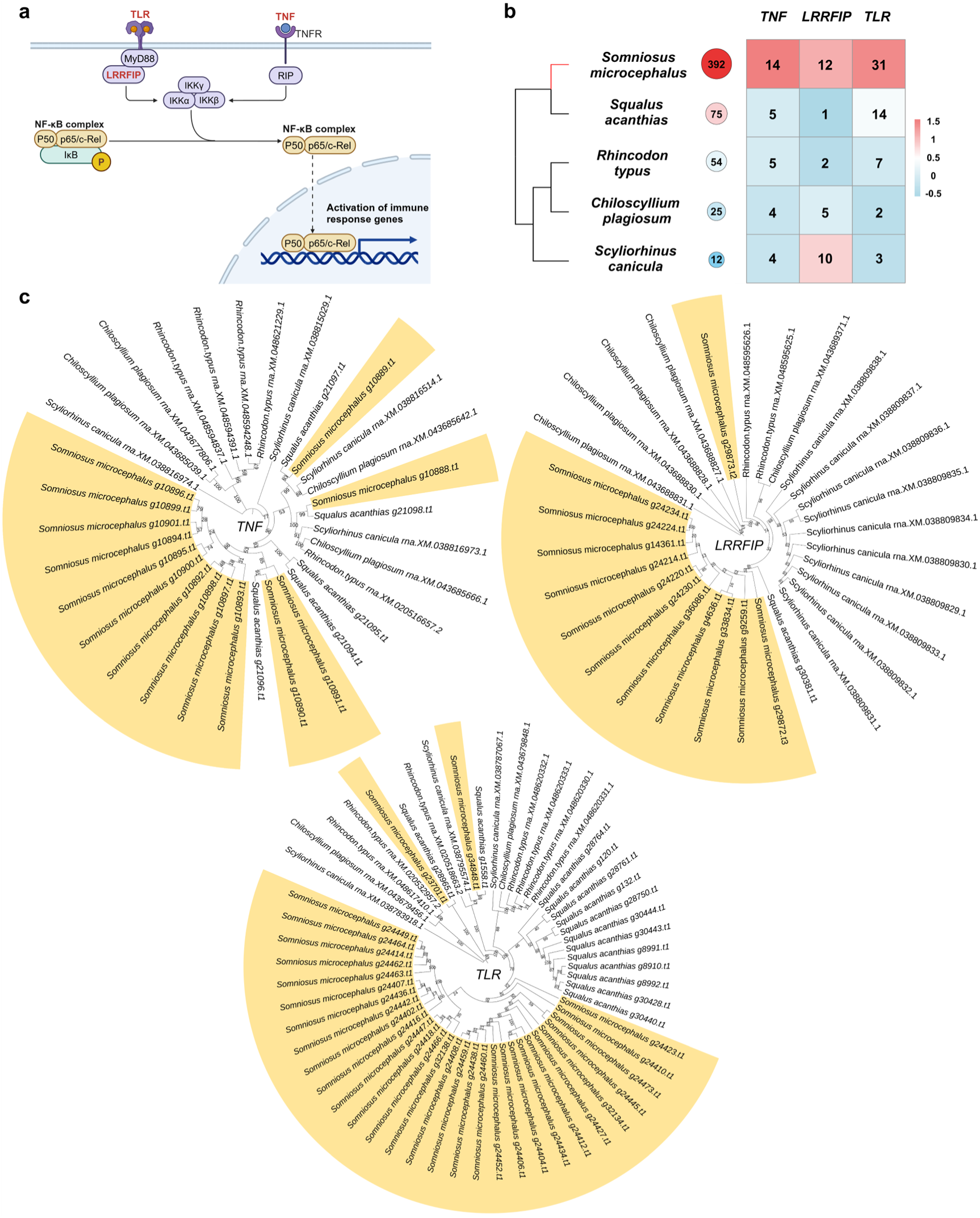
Analysis of NF-κB signaling pathway-related genes. (A) Functional diagram illustrating the involvement of genes from expanded gene families (labeled in red) in the NF-κB pathway in the *S. microcephalus* genome. (B) Comparison of the copy numbers of three genes among *S. canicula, C. plagiosum, R. typus, S. acanthias,* and *S. microcephalus* with documented lifespans. Numbers in the circles represent the maximum lifespan, while numbers in the heatmap indicate gene copy numbers. The value was calculated as the difference between the gene copy number of each species minus the average gene copy number of all five species divided by the standard deviation of the gene copy number of all five species. (C) Phylogenetic tree of three expanded genes involved in the NF-κB signaling pathway in five species.

### Positive selection of cancer-related genes as a potential key for long life

We calculated the rate ratio (ω) of non-synonymous (*K*_a_) to synonymous (*K*_s_) nucleotide substitutions in the coding regions of single-copy orthologous genes in various chondrichthyans. This analysis revealed 18 positively selected genes in *S. microcephalus*, which were then analyzed for GO and KEGG enrichment. We manually checked all 18 positively selected genes and summarized their potential functions relevant to the evolution of *S. microcephalus* adaptations in Table 2 and Supplementary files 4 &5. We identified several orthologs under positive selection in the *S. microcephalus* genome (*NOL6*^32^, *CTNNBL1*^33^, *UPK3B*^34^, *TMEM229A*^35^, *FOXF2*^36^, *FSCN1*^37^, *MAD2L1BP*^38^) that have been shown to be involved in cancer-related processes.

**Table 2.** Positively selected genes associated with longevity and adaptive trait.

| Function | Positive selected genes |
| --- | --- |
| Cancer | *NOL6, CTNNBL1, UPK3B, TMEM229A,*<br>*\*FOXF2, \*FSCN1, \*MAD2L1BP* |
| Vision | *POU4F1, GUCA1A, VPS11* |

\*Marker genes for cancer

GO enrichment analysis revealed significant enrichment in terms related to cell cycle regulation, proliferation, and differentiation, such as regulation of cell cycle, negative regulation of nuclear division, and negative regulation of cell division (Fig. 4A). KEGG enrichment analysis identified several pathways, including phototransduction, glutathione metabolism, ribosome biogenesis in eukaryotes, and spliceosome (Fig. 4B). Ten of the proteins encoded by the 18 positively selected genes are involved in protein-protein interactions and are divided into four smaller networks (Fig. 4C).

**Figure 4.**
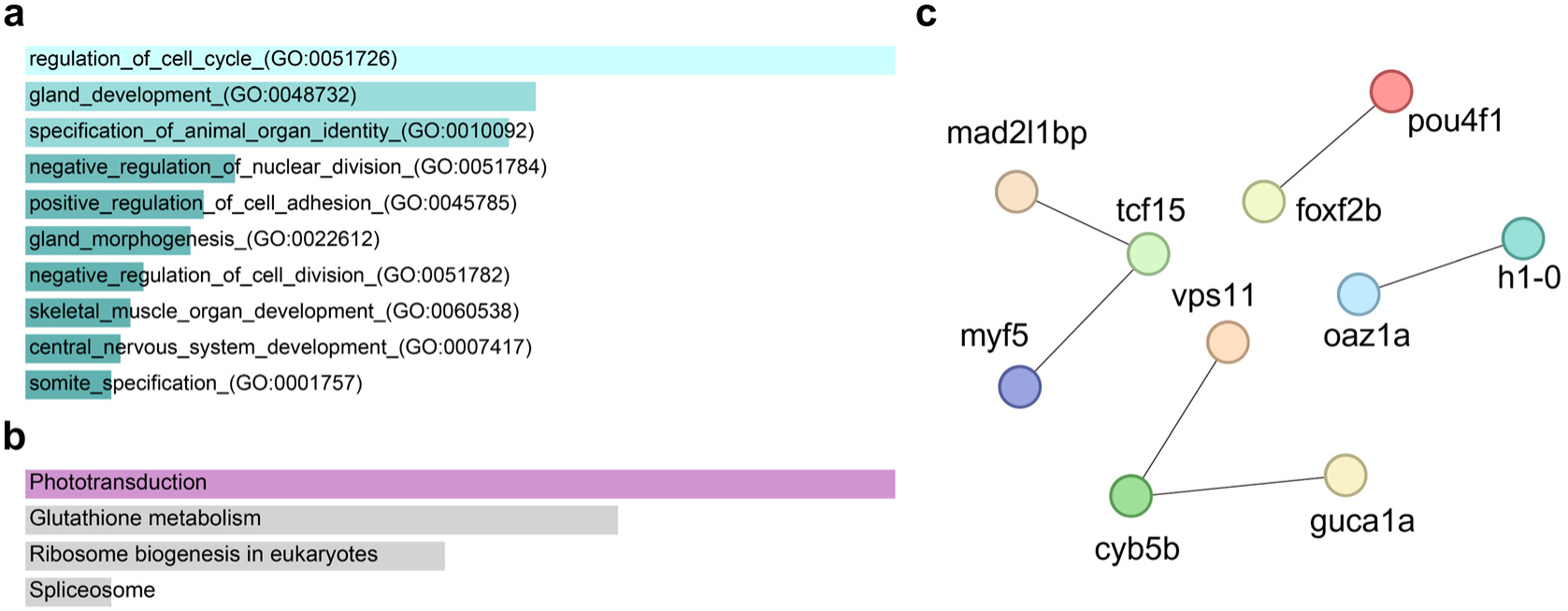
Summary of the functions of positively selected genes. (A) Top ten GO biological process terms for positively selected genes. (B) Top four KEGG enrichment terms for positively selected genes. (C) Protein-protein interaction network of positively selected genes.

### Dim-light vision suggested by the sequence signature of the photoreceptor

We identified partial segments of the rhodopsin (*RHO*) gene in the genome of *S. microcephalus*. First, RHO protein sequence alignment was performed between *S. microcephalus* and *R. typus*. The causative amino acid residues of RHO in *R. typus* are located at sites 94 and 178, with unique substitutions at these sites leading to reduced thermal stability of rhodopsin, which is the molecular basis for the human disease congenital stationary night blindness (CSNB)^39^. Our alignment confirmed that *S. microcephalus* does not possess these substitutions.

We then expanded the alignment to include more species. We found that the amino acid sequence of *S. microcephalus* RHO was identical to that of *G. melastomus*, a typical deep-sea dweller, at all eight amino acid residues (Table 3). These were previously characterized as spectral tuning sites, which affect RHO absorption wavelength^40^.

**Table 3.** Comparison of sites known to influence absorption wavelengths in rhodopsin.

| Species | $\lambda(\text{nm})^*$ | The residue numbers based on bovine RHO | | | | | | | | | |
| --- | --- | --- | --- | --- | --- | --- | --- | --- | --- | --- | --- |
|  |  | 83 | 94 | 122 | 124 | 132 | 178 | 208 | 261 | 292 | 299 |
| *S. microcephalus* | - | N | T | E | S | A | Y | F | F | S | A |
| *C. punctatum* | 500 | D | T | E | S | A | Y | F | F | A | A |
| *R. typus* | 478 | D | A | E | S | A | F | F | F | A | A |
| *S. canicula* | 494 | D | T | E | G | A | Y | F | F | A | S |
| *G. melastomus* | 483 | N | T | E | S | A | Y | F | F | S | A |
| *C. amblyrhynchos* | 504 | D | T | E | A | A | Y | F | F | A | S |
| *C. melanopterus* | 505 | D | T | E | A | A | Y | F | F | A | S |
| *L. erinacea* | 498 | D | T | E | G | A | Y | F | F | A | A |
| *S. torazame* | 484 | D | T | E | S | A | Y | F | F | S | S |

\*The wavelength of maximum absorption ( $\lambda$ max)

### Divergence and demographic history of the *Somniosus* species

Using the newly constructed genome assembly of *S. microcephalus* as a reference, we conducted population genetic analyses of two *Somniosus* species by mapping HiFi reads of *S. microcephalus* and newly sequenced shotgun sequencing data from a single *S. pacificus* individual.

PSMC analysis revealed a declining trend in the effective population size (*N_e_*) of *S. microcephalus* from the past through to the present, with the most recent estimate during the period 240-505 Kya being 1.55×10^4^ (Fig. 5A). In contrast, the *N_e_* of *S. pacificus* demonstrated signs of recovery following a historical bottleneck (Fig. 5B).

**Figure 5.**
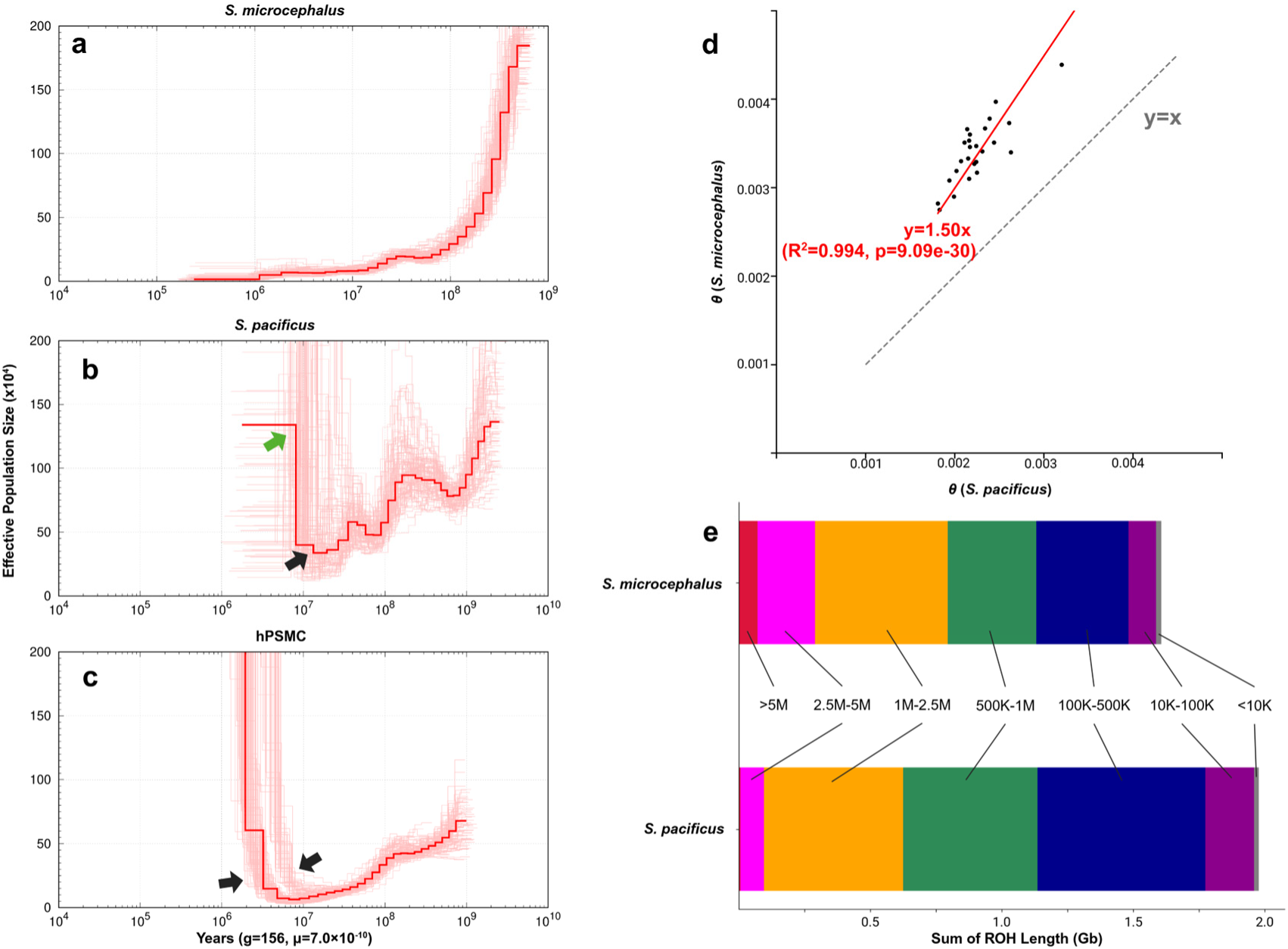
Population genetics of *S. microcephalus* and *S. pacificus*. (A) PSMC analysis for *S. microcephalus*. (B) PSMC analysis for *S. pacificus*. Black arrow: bottleneck, Green arrow: population expansion. (C) hPSMC analysis for the two *Somniosus* species. The arrow indicates the approximate timing of the onset of genetic isolation. (D) Comparison of heterozygosity between the two species. Each point represents a scaffold. The red line indicates the regression line. (E) The total base length of Runs of Homozygosity (ROH) is color-coded by the length of each ROH.

The hPSMC method^41^ simulates a hybrid diploid genome from two closely related populations and detects the divergence in estimated *N_e_* through PSMC analysis to estimate the divergence time between the two populations. In our analysis, the two species showed genetic isolation signals of approximately 10 Ma and 3 Ma (Fig. 5C).

We estimated the heterozygosity (θ) for scaffolds containing over 10 Mb of shared high-quality call sites in both *S. microcephalus* and *S. pacificus* (Fig. 5D). The long-term effective population size (*N_e_*), derived from θ≈4 *N_e_*μ, was estimated to be 9.82×10^5^∼1.95×10^6^ (2.75×10^-3^≦θ≦5.47×10^-3^) for *S. microcephalus*, and 6.46×10^5^∼1.44×10^6^ (1.81×10^-3^≦θ≦4.04×10^-3^) for *S. pacificus*. The θ values for the two individuals exhibited a consistent ratio across all scaffolds, precisely regressing to θ (*S. microcephalus*) =1.50θ (*S. pacificus*) (Fig. 5D).

To eliminate marker density-dependent bias, Runs of Homozygosity (ROH) were detected using high-quality shared call sites between the two sleeper shark species. The total base length of ROH was longer in *S. pacificus*, measuring 1.98 Gb, compared to 1.60 Gb in *S. microcephalus* (Fig. 5E). The genome of *S. microcephalus* contained a relatively higher abundance of long ROHs exceeding 2.5 Mb, resulting in a genomic inbreeding coefficient (F_ROH_)^42^ of 0.049, whereas *S. pacificus* had few long ROHs, with a F_ROH_ of 0.016.

## Discussion

Our study provides resources for the field of chondrichthyan genomics and insights into the evolution of exceptional longevity in nature. While *S. microcephalus* has garnered significant attention for its remarkably long lifespan^15^, its complete genome has not previously been sequenced. This likely relates to the challenges of dealing with the large genome size characteristic of elasmobranchs^43^. Here, we assembled the chromosome-level genome of *S. microcephalus* for the first time. Even with our assembly size reaching 5.9 Gb, the high-resolution chromatin interaction signals in our Hi-C data (Fig. 1B) demonstrate that our assembly has high accuracy, with nearly 90% of the contigs attached to 42 pseudochromosomes. Multifaceted assessments across the four assemblies revealed that our assembly surpasses those of other species in contiguity, achieving the highest N50 value, while also attaining the best score in completeness.

*S. microcephalus* possesses the largest genome size and the highest proportion of repetitive sequences among the chondrichthyan species studied to date, primarily due to transposons in this taxon rather than polyploidization or demi-duplication events^44^.

We have shown that the three chondrichthyan species, including *S. microcephalus*, share an ancient whole-genome duplication event from the era before their divergence, suggesting that modern elasmobranchs have maintained the ploidy from their common ancestor. A study on plants indicated that genome damage is not necessarily due to inefficient DNA repair in larger-genome species^45^. Research on the white shark further suggests that organisms with larger genomes might have enhanced DNA damage response and repair mechanisms to cope with increased DNA damage risk^20^. Consistently, our GO enrichment analysis of *S. microcephalus* shows that expanded gene families contribute to chromosome stability (Fig. 2B), suggesting a parallel evolution of this trait to prevent aging and DNA damage-related diseases.

KEGG enrichment analysis revealed numerous disease-related pathways (Fig. 2C), indicating *S. microcephalus* has a robust immune defense, potentially reducing the risk of infectious diseases and autoimmune issues. Previous studies have identified a unique lysozyme, β-N-acetylglucosaminidase, in the Leydig organs and ovaries of *S. microcephalus*, which may combat bacteria and parasites^46^. These findings further support the idea that *S. microcephalus* possesses a strong immune defense mechanism.

Theoretically, large, long-lived animals face a higher cancer risk due to more cells and cell divisions, yet cancer incidence is not as high as expected, known as Peto’s paradox^47^. Three genes identified as positively selected in our study *FOXF2, FSCN1,* and *MAD2L1BP* are considered biomarkers for cancer (Table 2). *FOXF2*, a crucial forkhead box transcription factor, regulates the tumor immune microenvironment. Its aberrant expression affects the epithelial-mesenchymal transition (EMT), G1-S cell cycle transition, and Wnt/β-catenin signaling, impacting tumor development^36^. *FSCN1* encodes an actin-binding protein involved in cell migration, motility, and adhesion. Previous studies show it is a critical downstream component in pathways associated with tumor progression^37^. *MAD2L1BP*, encoding p31comet, is involved in processes such as spindle assembly checkpoint silencing, accurate chromosome segregation, and homology-directed DNA repair. It is hypothesized to be a potential cancer treatment target^38^. These genes may provide insights into how *S. microcephalus* mitigates cancer risk, addressing Peto’s paradox.

We propose that the classical NF-κB signaling pathway is crucial for the exceptional longevity of *S. microcephalus*, as it regulates cell proliferation, migration, DNA repair, apoptosis, and immune response, which are all closely linked to inflammation, cancer, and autoimmune diseases^48^. *TNF*, one of the most effective NF-κB pathway activators, plays a broad role in inflammation, immune regulation, tumor lysis, and apoptosis^49^. The *TLR* gene family, encoded pattern recognition receptors that vary in number and type among species, primarily functions in the innate immune system^50^. A recent genomic analysis of the long-lived red sea urchin identified a high copy number of *TLR* genes, highlighting their importance in the immune system^51^. In the canonical NF-κB pathway, LRRFIP influences NF-κB activation by interacting with MyD88 and participating in the TLR pathway^48,52^. Our results indicate that the *S. microcephalus* genome has significantly higher copy numbers of the three gene families compared to four other relatively short-lived shark species (see Fig. 3B), suggesting stronger immune functions and enhanced cellular protection and repair capabilities. Additionally, studies show that dysregulation of NF-κB can lead to continuous tumor cell proliferation, prevention of apoptosis, and evasion of host immune system attacks^48^. NF-κB exhibits complex mechanisms in cancer development, potentially promoting or inhibiting tumor growth and inducing or suppressing apoptosis, depending on the organ context^53^. Therefore, *S. microcephalus* may have acquired refined regulatory mechanisms for moderate NF-κB activation, avoiding the negative effects of its continuous activation.

We identified 14 copies of the *CGNL1* gene in *S. microcephalus* (Supplementary File 2), encoding a protein localized at adherens junctions and tight junctions. The *CGNL1* gene was positively selected in both the short-lived turquoise killifish and the long-lived Brandt’s bat^54^. Furthermore, *CGNL1* appeared in a single nucleotide polymorphism model based on a genome-wide association study of 801 centenarians^55^. Our findings confirm the potential of the *CGNL1* gene as a target for longevity studies.

The maximum absorption wavelength (λ_max_) of RHO shifts towards 480 nm with increasing habitat depth^56^. All eight RHO sites in *S. microcephalus* align with those of *G. melastomus* (Table 3), suggesting a likely blue shift in *S. microcephalus* RHO. Moreover, during field experiments in Svalbard, it was observed that *S. microcephalus* responded clearly to submersible light, or nearby movements when animals were on deck, indicating underestimated visual capacity (attributed to copepod parasite attachment). Newborns likely experience a parasite-free period, and studies have shown that parasite attachment is not permanent; scars on the cornea indicate previous attachment points^57^. Though *S. microcephalus* ecology across growth stages has not been sufficiently studied, these parasite-free periods may have influenced gene evolution.

*S. microcephalus* lives in dark environments much of its life and likely relies heavily on olfaction. A study on its olfactory rosette structure found a larger lamellar surface area than in other species, indicating a superior sense of smell^58^. Additionally, brain morphology comparisons reveal a smaller optic tectum and an enlarged olfactory region, highlighting olfactory dependence in *S. microcephalus*^59^. Previous studies have shown that *C. milii* and three other shark species exhibit a highly degenerated olfactory receptor gene repertoire^44,60^. Our analysis did not detect complete olfactory receptor genes in *S. microcephalus*. However, 454 species-specific gene families showed significant enrichment in the olfactory transduction pathway (Fig. S3), particularly around the “*K02A2.6-like*” gene, which remains unstudied. We suggest that these species may use non-traditional molecular mechanisms for olfaction, warranting further research into these unidentified genes.

Large elasmobranchs, including the genus *Somniosus*, face high extinction risks due to their slow maturation rates^61^, which necessitates special attention from a conservation standpoint. Utilizing the newly constructed *S. microcephalus* genome assembly, we conducted the first genome-wide population genetic analysis for the genus *Somniosus*. Notably, the PSMC patterns for *S. pacificus* were unstable across bootstrap replicates and differed from *S. microcephalus* even in older periods when they were presumably part of the same population (Fig. 5A, B). This discrepancy stems from using the *S. microcephalus* genome as a reference for *S. pacificus* and the low sequencing depth for *S. pacificus*. Such limitations suggest that while PSMC can accurately predict general *N_e_* trends, absolute *N_e_* values, and time estimates remain less reliable^62^. It is reasonable to infer that *S. pacificus* followed the same *N_e_* trajectory as *S. microcephalus* until they diverged, after which it experienced a bottleneck and expansion (black and green arrows, Fig. 5B). The hPSMC analysis indicated two signals of population divergence (Fig. 5C), suggesting three ancestral populations formed via a two-step divergence. The estimated complete divergence around 3 Ma aligns with previous studies (1-3.5 Ma)^29,30^.

Heterozygosity regressed precisely to the equation θ (*S. microcephalus*) =1.50θ (*S. pacificus*) across scaffolds with shared sites (Fig. 5D). Given that θ≈4*N_e_*μ, and assuming identical mutation rates (μ) for these closely related species, it can be inferred that the long-term *N_e_* of *S. microcephalus* is approximately 1.5 times that of *S. pacificus*. Despite the recent *N_e_* recovery seen in *S. pacificus* (green arrow, Fig. 5B), the lower θ indicates a significant bottleneck before its expansion (black arrow, Fig. 5B).

*S. microcephalus* exhibited a higher abundance of long ROHs compared to *S. pacificus* (Fig. 5E). The F_ROH_ related to long ROHs is known to correlate with the pedigree inbreeding coefficient^42^, suggesting more recent inbreeding in *S. microcephalus*, consistent with the PSMC *N_e_* dynamics (Fig. 5A, B). Conversely, *S. pacificus* showed a higher base length of fragmented ROHs, suggesting historical inbreeding, aligning with the bottleneck hypothesis derived from heterozygosity analysis. Our synthesis of three population genetic analyses coherently clarifies the *N_e_* dynamics in these species.

Future research should involve sequencing multiple individuals from various marine regions to elucidate global population dynamics in greater detail. Given *Somniosus*’s long generation times^15^ and low mutation rates^63^, recent resource fluctuations may be hard to detect through PSMC and heterozygosity analyses.

However, ROH analysis could identify recent inbreeding events, providing insights into short-term dynamics. Nevertheless, the genome assembly we have constructed will undoubtedly serve as a crucial reference sequence for future studies.

## Methods

### Sample collection and nucleic acid extraction

In 2021, a female *S. microcephalus* (total length: 345cm) was captured in Kongsfjorden, Svalbard Archipelago, for ongoing biotelemetry studies^12,64^. A fin clip and blood samples were collected prior to release of the animal. These tissues were preserved in a DNA/RNA shield (ZYMO RESEARCH), and rapidly frozen for subsequent analysis.

Genomic DNA was extracted from 60 mg of the sampled *S. microcephalus* fin using the Nucleobond HMW DNA kit (MACHEREY-NAGEL) as per the manufacturer’s protocol. The DNA solution’s concentration and size distribution were verified using Tapestation (Agilent) and Nanodrop (Thermo Fisher Scientific). Total RNA was extracted from 5 mg of frozen fin sample and 200 µl of frozen blood using the ReliaPrep RNA Tissue Miniprep System kit (Promega) and the Nucleospin RNA Blood kit (MACHEREY-NAGEL) following the manufacturer’s guidelines.

### Library construction and sequencing

For HiFi read generation, the PacBio Single-Molecule Real-Time (SMRT) libraries were prepared using the SMRTbell Express Template Prep Kit 2.0 and SMRTbell prep kit 3.0 (Pacific Biosciences) and sequenced in CCS mode on the PacBio Sequel II and PacBio Revio systems. Subsequently, circular consensus analysis using subreads was conducted with SMRT Link v10.0 and Google Health DeepConsensus, which yielded 229.7 Gb HiFi reads.

For Hi-C library preparation, 100 mg of the frozen fin sample immersed in RNAlater was utilized. The Hi-C library was prepared according to the iconHi-C protocol^65^. Employing restriction enzymes DpnII and HinfI, it was sequenced on a HiSeq X sequencing platform to obtain 395.4 Gb Hi-C read pairs (Illumina Inc., CA, USA).

### Transcriptome sequencing

For transcriptome sequencing, the libraries were constructed from Total RNA extracted from fin and blood using the Illumina Stranded mRNA Prep Ligation (Illumina). The PCR cycle for library amplification was set to 11 cycles. Quality checks and concentration measurements of the prepared libraries were performed using Tapestation. Pair-end sequencing was conducted on the Illumina HiSeq X platform.

### Genome assembly and species identification

Initially, the HiFi reads generated by the PacBio platform were assembled using Hifiasm (v0.19.5-r587)^66^ with the standard default parameters. Potential sequencing errors were corrected through all-vs-all sequence alignment. Based on sequence overlaps, a phased string graph was constructed, resulting in contig generation for the next assembly step. Potential haploids and contig overlaps in the *de novo* assembly were removed using Purge_Dups (v1.2.5)^67^ step by step. Coverage-related calculations and self-alignments were conducted using Minimap2 (v2.26-r1175)^68^ with HiFi reads.

The paired-end Hi-C data were used to scaffold the primary assemblies obtained from Hifiasm. Specifically, Hi-C reads were first quality trimmed using Trim Galore (v0.6.10, https://www.bioinformatics.babraham.ac.uk/projects/trim_galore/), and then mapped to the assembled genome via Chromap (v0.2.5-r473)^69^. The Hi-C scaffolding tool YaHS (v1.2a.1)^70^ was used to create a contact matrix, construct and prune a scaffolding graph, and finally output scaffolds. PCR/optical duplicates were marked by biobambam2 (v2.0.183, http://www.sanger.ac.uk/science/tools/biobambam) as suggested by YaHS’s author. The Hi-C contact map was generated using Pre provided by YaHS^70^ and Juicer Tools (v1.19.02)^71^ and visualized using Juicebox Assembly Tools (v2.17.00)^72^. The final genome assembly was built to be consistent with the chromatin contact profiles.

The quality of the genome assembly was evaluated using various software tools: Genome completeness was assessed via compleasm^73^ in the genome model based on the Vertebrata dataset. Genome size, N50 length, fragment count, and GC content were calculated using QUAST (v5.2.0)^74^. Consensus quality value (QV) and k-mer completeness were calculated using Merqury (v1.3)^28^, and coverage was determined by samtools (v1.19.2)^75^. Finally, the assembly statistics were compared to other shark species.

Sequences from the cytochrome b region (mitochondrial genome) and the *RAG1* and *ITS2* regions (nuclear genome) of *S. microcephalus* published on NCBI were used as queries to perform a BLASTN (v2.14.0+)^76^ homology search against the obtained genome assembly to identify the corresponding sequences for this individual. For the nuclear genome region in the genome assembly, read-level validation was performed, considering potential sequence deletions in one of the parents.

### Genome annotation

Repeat sequence annotation of the *S. microcephalus* genome was conducted prior to gene model prediction. Initially, we accessed the integrated Dfam (v3.7)^77^ and Repbase_20181026^78^ database using famdb.py of RepeatMasker (v4.1.5)^79^ to export the repeat families of ‘Squaliformes’ and its ancestor nodes along with all subsequent taxa, as a homology-based database. For *de novo* prediction, a library of ab initio repeat sequences for the *S. microcephalus* genome was constructed using RepeatModeler (v2.0.4)^80^. Subsequently, we combined both the homology-based and *de novo* databases and used RepeatMasker^79^ for soft-masking of the repetitive sequences. Tantan (v49)^81^ was used to further mask simple tandem repeats that were not detected by RepeatMasker.

The soft-masked genome was used for protein-coding gene prediction using BRAKER3 (v3.0.7)^75,82–90^. RNA-seq data obtained from fin tissue and blood were first quality trimmed using Trim_Galore, then mapped to the soft-masked genome using HISAT2 (v2.2.1)^91^ as RNA-seq evidence input file for the BRAKER3 pipeline, specifying the parameters ‘verbosity=4’. The output GFF file of BRAKER3 was processed using the scripts agat_convert_sp_gxf2gxf.pl, agat_sp_fix_overlaping_genes.pl, agat_sp_add_start_and_stop.pl, and agat_sp_filter_incomplete_gene_coding_models.pl in the Another Gtf/Gff Analysis Toolkit (AGAT, v1.2.0)^92^ to clean up the gene model, correct overlapping predictions, and remove or correct protein-coding genes with CDS less than 100 amino acids and incomplete genes. OMAmer^93^ and OMArk^94^ were utilized to measure the completeness of the proteome and identify the presence of potential contamination from other species. After removing potential contamination, the same scripts in the AGAT^92^ were used again to obtain final gene model predictions. The completeness of the predicted gene models was assessed using compleasm^73^ in the protein model. For functional annotation, the integrated gene sets were queried against the NR database (ftp://ftp.ncbi.nlm.nih.gov/blast/db/) with diamond (v2.1.8)^87^. Additionally, InterProScan (v5.59_91.0)^95^, eggNOG-mapper Web^96,97^, and PANNZER^98^ were used for functional annotation based on multiple databases. Non-coding RNA (ncRNA) was annotated using Infernal (v1.1.5)^99^ based on the Rfam (v14.10)^100^ database.

### Evolutionary analysis of gene families

To characterize the expansion and contraction of gene families in *S. microcephalus* during evolution, we downloaded 12 high-quality chondrichthyan genomic datasets from NCBI, including *A. radiata, L. erinacea, P. pectinata, M. hypostoma, H. sabinus, S. acanthias, H. ocellatum, C. plagiosum, R. typus, S. tigrinum, S. canicula, C. milii,* and selected *D. rerio* as an outgroup (Table S5). Genes of the 14 species were clustered using OrthoFinder (v2.5.5)^31^ with default parameters to identify orthologs and paralogs. Protein sequences of single-copy orthologs from each species were aligned and translated into corresponding coding DNA sequences using MUSCLE (v3.8.1551)^101^ and PAL2NAL (v14.1)^102^. Poorly aligned positions were filtered out with Gblocks (v0.91b)^103^. The best model ‘GTR+I+G4’ was selected by modeltest-ng (v0.1.6)^104,105^, followed by 1000 bootstrap replicates executed with RAxML (v8.2.13)^106^ to generate the phylogenetic tree. The ultrametric tree was generated by MCMCtree from the PAML (v4.10.7)^107^ package, based on the species divergence times (*D. rerio* and *C. milii* diverged at 440.8-495.2 MYA, *S. microcephalus* and *S. acanthias* diverged at 137.3-170.0 MYA) obtained from TimeTree^108^. Gene families with excessive copy number differences between species were filtered out and subsequently analyzed using CAFE (v5.1.0)^109^ with the Gamma model and Poisson distribution option to determine the number of gene family expansions and contractions on each evolutionary branch. Genes with significant expansions and contractions in *S. microcephalus* were extracted and analyzed for Gene Ontology (GO) and Kyoto Encyclopedia of Genes and Genomes (KEGG) enrichment using clusterProfiler (v4.10.1)^110^, with a P-value threshold of 0.05. The NF-κB signaling pathway-related gene trees were constructed similarly to the species tree, with species lifespan data obtained from AnAge^111^.

To calculate the synonymous substitutions per site (*K*_s_) distribution for the four species *S. microcephalus*, *C. plagiosum*, *S. canicula*, and *D. rerio*, we employed the following methodology - protein sequences were self-aligned using DIAMOND^87^, followed by the identification of paralogs in the syntenic regions using MCScanX^112^. *K*_s_ values were then calculated using ParaAT.pl^113^ and KaKs_Calculator2.0^114^, and subsequently visualized.

### Positive gene selection identification

Positively selected genes in the *S. microcephalus* genome were identified by quantifying synonymous substitutions (*K*_s_) and non-synonymous substitutions (*K*_a_). Screening of positively selected genes exhibiting the *K*_a_/*K*_s_ value of larger than 1 was performed using single-copy orthologous genes based on the result of OrthoFinder between *S. microcephalus* and related species, along with the phylogenetic tree generated by RAxML^106^. The branch-site model (model = 2; NSsite = 2) of codeml from the PAML (v4.10.7)^107^ package was used for the positive selection analysis, with *S. microcephalus* set as the foreground branch. Specifically, the ‘cleandata’ option of codeml was enabled, setting ‘fix_omega = 1; omega = 1’ for the null model and ‘fix_omega = 0; omega = 2’ for the alternative model. The Chi2 program in PAML^107^ was then used to check and correct for multiple hypotheses (p-value ≤ 0.05). The posterior probability of positively selected sites was calculated using the Bayes Empirical Bayes (BEB) method. Sites with a probability value ≥ 0.9 indicated that the gene was under positive selection on the target branch. We then manually inspected the alignment of all positively selected gene protein sequences and removed potential false-positive results. Positively selected genes with gene ID of *D. rerio* were subsequently analyzed for GO and KEGG enrichment using FishEnrichr^115,116^. The protein-protein interaction network of proteins encoded by positively selected genes was constructed using STRING^117^.

### Rhodopsin sequence comparison

The rhodopsin (RHO) sequence of the *R. typus* (XP_048462427.1) downloaded from NCBI was used as queries to conduct a similarity search with TBLASTN (v 2.14.0+)^118^ against the genome assembly of *S. microcephalus* to obtain the RHO ortholog. Multiple sequence alignments of the obtained RHO sequences and the published RHO sequences of other chondrichthyan species: *Scyliorhinus torazame*^44^, *Carcharhinus amblyrhynchos* (QGW08842.1), *Carcharhinus melanopterus* (QGW08844.1), *Chiloscyllium punctatum* (QGW08841.1), *Leucoraja erinacea* (XP_055503835.1), *Scyliorhinus canicula* (XP_038666779.1) and *Galeus melastomus* (O93441.1) were performed using MAFFT^119^ to manually check for eight sites thought to affect the absorption spectra of RHO.

### Genome sequencing of *S. pacificus*

To understand comparative population dynamics, specimens of the Pacific sleeper shark *S. pacificus* were collected from Suruga Bay, Japan, in May 2019 and subsequently preserved at −4°C. DNA was extracted from the skeletal muscle of the trunk region using the DNeasy Blood & Tissue Kit (Qiagen). The extracted DNA was fragmented using the NEBNext dsDNA Fragmentase (New England Biolabs) and then prepared for sequencing with the Nextera DNA Library Preparation Kit (Illumina). Sequencing was performed on the Illumina HiSeq X platform.

### Population dynamics assessment

The final genome assembly of *S. microcephalus* was subjected to hard masking using RepeatModeler (v2.0.4)^80^ and RepeatMasker (v4.1.5)^79^. HiFi reads of *S. microcephalus* were mapped to the repeat-masked *S. microcephalus* genome assembly using minimap2 (v2.26-r1175)^68^, while 149 Gb of paired-end 150 bp shotgun sequencing data of *S. pacificus* were aligned using bwa (v0.7.17-r1198-dirty)^120^. The resulting alignments were converted to BAM files with samtools (version 1.4.1)^75^.

The average depth of unmasked regions was calculated per scaffold using the “samtools depth” command, and only BAM files corresponding to scaffolds with an average coverage of at least 25x^62^ were extracted. Variants were called using commands “samtools mpileup -C50 -q20 -Q20”, “bcftools call -c”, and “vcfutils.pl vcf2fq -d 1/3*(average coverage) -D 2*(average coverage)”, producing output in FASTQ format. These were then converted to PSMCFA format using the “fq2psmcfa -q20” command implemented in PSMC^121^. The PSMC analysis was performed with parameters ‘-N25 -t15 -r5 -p “4+25*2+4+6”’, including 100 bootstrap replicates, and visualized assuming a mutation rate of 7.0×10^-10^ /(bp・generation)^63^ and a generation time of 156 years^15^.

To estimate the divergence time between *S. microcephalus* and *S. pacificus*, hPSMC^41^ was applied. Variants were called from the respective mappings using the command “samtools mpileup -s -q30 -Q60 -r”, and haplotypes were subsequently called using “pu2fa” from the Chrom-Compare (https://github.com/Paleogenomics/Chrom-Compare). Scaffolds common to both species were extracted, and a hybrid PSMCFA was generated using the “psmcfa_from_2_fastas.py -b10 -m5” command. PSMC analysis and visualization were performed using the previously described parameters.

For each of the two shark species, variants were called from the BAM files generated in the PSMC section using the command “samtools mpileup -C50 -q20 - Q20”. Only sites meeting the minimum coverage of 1/3*(average coverage) and maximum coverage of 2*(average coverage) were output. The base frequencies at each site were converted into PRO files for each scaffold using bam2pro.awk implemented in mlRho (v2.9)^122^. From these PRO files, sites called in both species were extracted. The formatting was performed using FormatPro (v0.5) for scaffolds containing common sites totaling over 10 Mbp. MlRho was then used to estimate θ.

The θ values for each scaffold were visualized using Matplotlib in Python (v3.9.16), and a regression line with an intercept of 0 was calculated using statsmodels (https://www.statsmodels.org/stable/api.html).

Using the BAM files generated in the PSMC section, base frequencies were output in VCF format utilizing bcftools (v1.17)^75^ with the commands “bcftools mpileup -C50 -q20 -Q20 -Ou -a DP, AD -A” and “bcftools call -m -A -Oz”. Sites commonly called in both species were then extracted. Runs of homozygosity (ROH) were predicted using the command “bcftools roh -G30 --AF-dflt 0.4”^123^. The proportion of the genome occupied by ROH regions larger than 2.5 Mb was calculated as the genomic inbreeding coefficient (F_ROH_)^42^.

## Supporting information

Complete enrichment results for the unique gene family of the Greenland shark

Complete results for expansion and contraction of the Greenland shark gene family

Complete enrichment results for expansion and contraction of the Greenland shark gene family

Complete statistics of positively selected genes in Greenland shark

Complete enrichment results for positively selected genes in Greenland shark

## Acknowledgments

The authors thank Eric Ste Marie for assistance with fieldwork and the OceanXplorer crew and onboard scientists for providing a platform to undertake the sampling of the Greenland shark. We also thank Yasuda Satoshi and Koichi Shibukawa for their help in sampling Pacific sleeper shark tissues. This work was supported by JSPS KAKENHI Grant Number 20H00427.

## Data availability

All raw sequencing data (Hi-C reads, PacBio HiFi reads, and Illumina short reads) and final assembly can be found on NCBI under BioProject PRJNA1218902.

## Author contributions

S.K. conceived the study. S.A. and S.Ku. supervised the study. Y.Y.W., N.E.H., K.M.K., and C.L. provided materials. K.N. and M.K. performed the experiments. K.Y. and K.Yo. conducted genome assembly and annotation. K.Y. and H.M. performed the data analysis. K.Y. wrote the manuscript, with significant contributions from S.Ku., H.M., N.E.H., K.M.K., and C.L., as well as input from all authors. All authors read and approved the final manuscript.

**Correspondence** and requests for materials should be addressed to Shigeharu Kinoshita.

---

**Figure S1.**
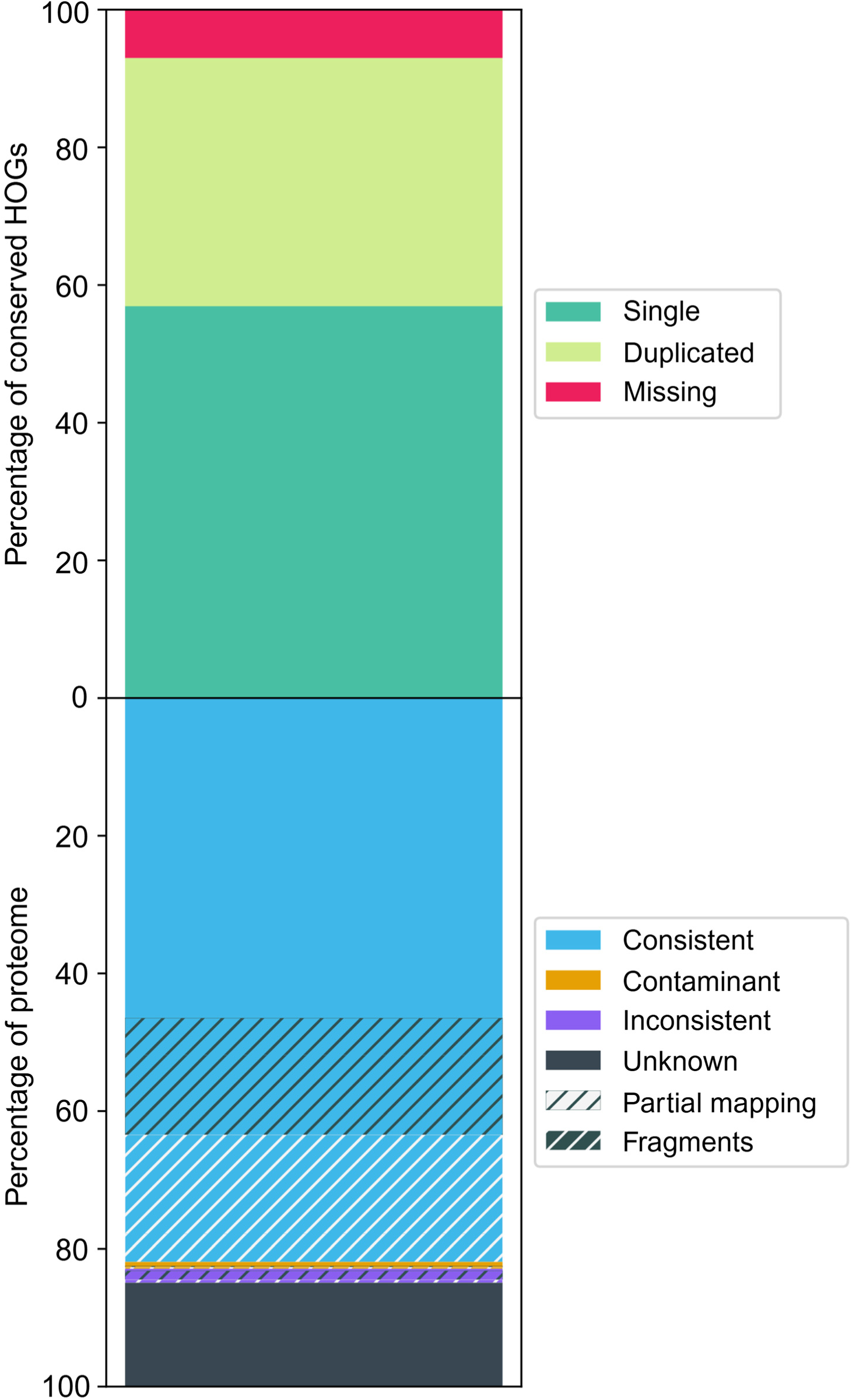
The completeness and whole proteome quality assessment by OMArk.

**Figure S2.**
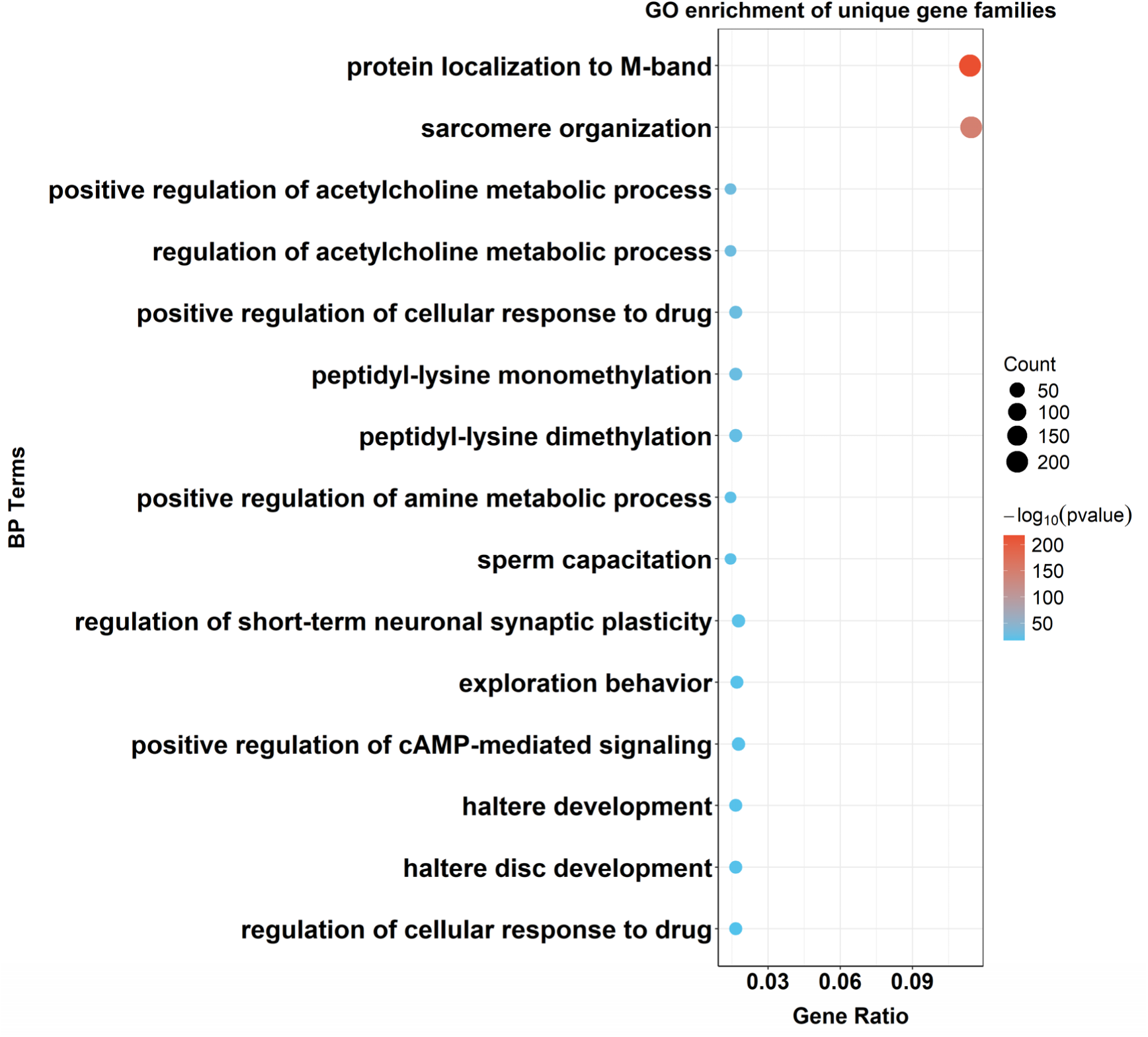
Top fifteen GO enrichment terms for unique gene families in *S. microcephalus*.

**Figure S3.**
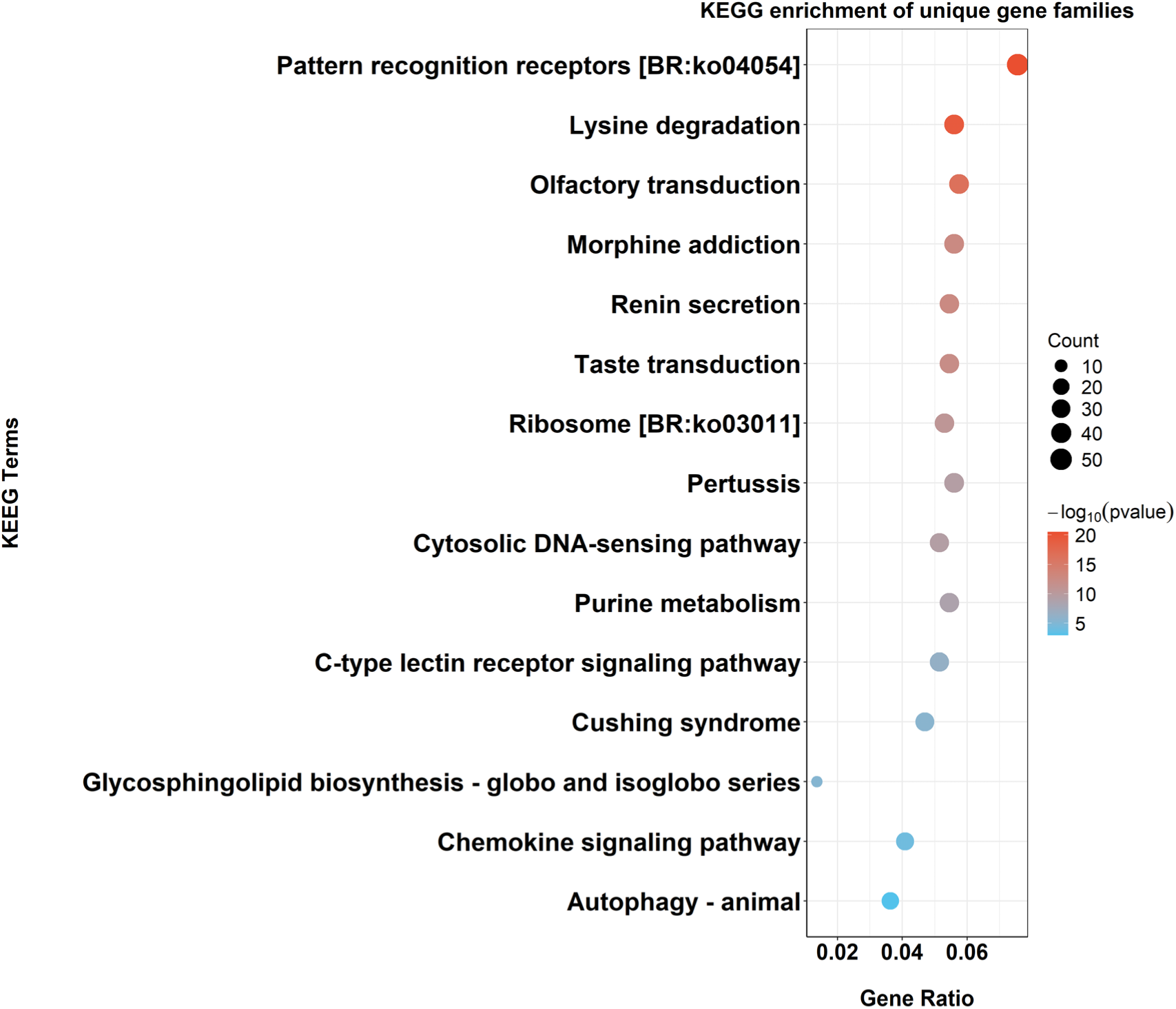
Top fifteen KEGG enrichment terms for unique gene families in *S. microcephalus*

**Table S1.** Repeat sequence classification result statistics.

| Type | length (bp) | Percent (%) |
| --- | --- | --- |
| DNA transposons | 230,512,066 | 3.88 |
| SINEs | 145,021,515 | 2.44 |
| LINEs | 1,500,564,655 | 25.25 |
| LTR elements | 1,111,409,632 | 18.70 |
| Tandem repeat | 767,082,845 | 12.90 |
| Total | 4,940,330,074 | 83.12 |

**Table S2.** Statistical analysis of protein-coding genes.

| Item | Number |
| --- | --- |
| Number of genes | 37,125 |
| Number of CDS | 44,055 |
| mean gene length (bp) | 30,700 |
| mean transcript length (bp) | 34,803 |
| mean CDS length (bp) | 1,275 |
| mean exon length (bp) | 200 |
| mean exons per CDS | 6.4 |
| Completeness score | 86.5% |

**Table S3.** Functional annotation of protein-coding genes for *S. microcephalus*.

| Type | Number | Percent% |
| --- | --- | --- |
| eggNOG | 26,774 | 72.0 |
| KEGG | 16,982 | 45.7 |
| UniProt | 25,351 | 68.2 |
| InterPro | 27,960 | 75.2 |
| Pfam | 24,514 | 65.9 |
| PANTHER | 27,217 | 73.2 |
| NR | 31,661 | 85.2 |
| GO | 19,572 | 52.6 |
| ALL | 32,132 | 86.5 |

**Table S4.** Statistical analysis of non-coding RNA.

| Item | Number |
| --- | --- |
| riboswitch | 7 |
| tRNA | 12,563 |
| rRNA | 906 |
| sRNA | 1 |
| snRNA | 1,221 |
| ribozyme | 26 |
| miRNA | 241 |
| frameshift_element | 2 |
| lncRNA | 9 |
| leader | 1 |
| antisense | 2 |
| IRES | 1 |
| Others | 1,198 |

**Table S5.** Statistical data for related species.

| Species | Num_scaffold | Size | N50 | Protein-coding | RefSeq |
| --- | --- | --- | --- | --- | --- |
| *Chiloscyllium plagiosum* | 101,976 | 3.6 Gb | 73.2 Mb | 19,916 | GCF_004010195.1 |
| *Rhincodon typus* | 16,776 | 2.9 Gb | 70.8 Mb | 20,753 | GCF_021869965.1 |
| *Pristis pectinata* | 172 | 2.3 Gb | 101.7 Mb | 17,434 | GCF_009764475.1 |
| *Hemiscyllium ocellatum* | 1,962 | 4 Gb | 83.6 Mb | 19,215 | GCF_020745735.1 |
| *Callorhinchus milii* | 1,761 | 991.5 Mb | 69.3 Mb | 17,192 | GCF_018977255.1 |
| *Scyliorhinus canicula* | 645 | 4.2 Gb | 198.8 Mb | 20,574 | GCF_902713615.1 |
| *Leucoraja erinacea* | 1,768 | 2.2 Gb | 57.2 Mb | 18,970 | GCF_028641065.1 |
| *Amblyraja radiata* | 957 | 2.6 Gb | 62.1 Mb | 18,474 | GCF_010909765.2 |
| *Hypanus sabinus* | 2,989 | 4 Gb | 112.3 Mb | 22,522 | GCF_030144855.1 |
| *Mobula hypostoma* | 200 | 3.5 Gb | 152.4 Mb | 18,916 | GCF_963921235.1 |
| *Squalus acanthias* | 3,882 | 3.7Gb | 124.1 Mb | 37,273 | GCA_030390025.1 |
| *Stegostoma tigrinum* | 1,266 | 3.8 Gb | 76.6 Mb | 18,281 | GCA_022316705.1 |
| *Danio rerio* | 1,917 | 1.4 Gb | 7.4 Mb | 26,448 | GCF_000002035.6 |

